# Following The Niche: Reconstructing 32,000 Years Of Niche Dynamics In Four European Ungulate Species

**DOI:** 10.1101/2020.12.07.401133

**Authors:** Michela Leonardi, Francesco Boschin, Paolo Boscato, Andrea Manica

## Abstract

An understanding of how ecological niches can change through time is key to predicting the effect of future global change. Past climatic fluctuations provide a natural experiment to assess the extent to which species can change their niche. Here we use an extensive archaeological database to formally test whether the niche of four European ungulates changed between 40 and 8 kya (i.e. before major anthropogenic habitat modification and excluding the confounding effect of domestication). We find that niche change depended on habitat. Horse and aurochs, which are adapted to open environment, changed their niche after the Last Glacial Maximum, and it is unclear whether this was the result of adaptation, or an expansion of the realized niche as a response to the extinction of other megafauna (competitors and predators) that shared the same habitat preferences. On the other hand, red deer and wild boar, which prefer close and semi-close habitats, did not change their niche during the same period; possibly because these habitats have experienced fewer extinctions. Irrespective of the mechanism that might have led to the observed niche changes, the fact that large mammals with long generation times can change their niche over the time period of thousands of years cautions against assuming a constant niche when predicting the future.

**Significance statement:** When predicting species responses to future change, it is often assumed that their habitat preferences (i.e. their niche) will not change. However, it is strongly debated whether this is reasonable. Here we show that two out of four species of large European ungulates changed their niche following the Last Glacial Maximum, possibly as a response to the reorganization of animal communities that resulted from numerous megafauna extinctions. This finding cautions against the assumption of a constant niche, highlighting that, to predict the future, we will ultimately need to understand the mechanisms that underpin the success of a given species under different climatic conditions.

## Introduction

Given the current rate of climatic changes due to human activity (1), there is an urgent need to establish the best possible framework to study how animal species react to climatic fluctuations and how their ecological niche may vary through time. The climate fluctuations that characterized the last tens of thousands of years can be used as a virtual lab to gain a better understanding of the ecological niche dynamics under different environmental conditions, which in turn may help defining better conservation strategies for the future (2).

The ecological niche of a species as defined by Hutchinson in 1957 (3) is defined in two different ways. The fundamental niche is the range of environmental conditions (intended as both abiotic variables and biological relationships) in which the species can survive and reproduce. However, certain conditions that would, in theory be suitable, might be realized in the physical space that can be occupied by the species at a given time; the subset of conditions that are available, and thus used by a species, are defined as the realized niche. In the present work we will only focus on the latter.

When dealing with anthropogenic climate change, the question becomes: will there be enough realized niche for the species to survive under the new climatic conditions? An answer can be found using Species Distribution Models (SDMs) (4). Under this framework, occurrences of a given species are associated with climatic variables of the area they inhabit (Figure 1), producing a model that, by identifying the realized niche, can predict its potential geographic distribution based on suitable climate.

**Figure 1:**
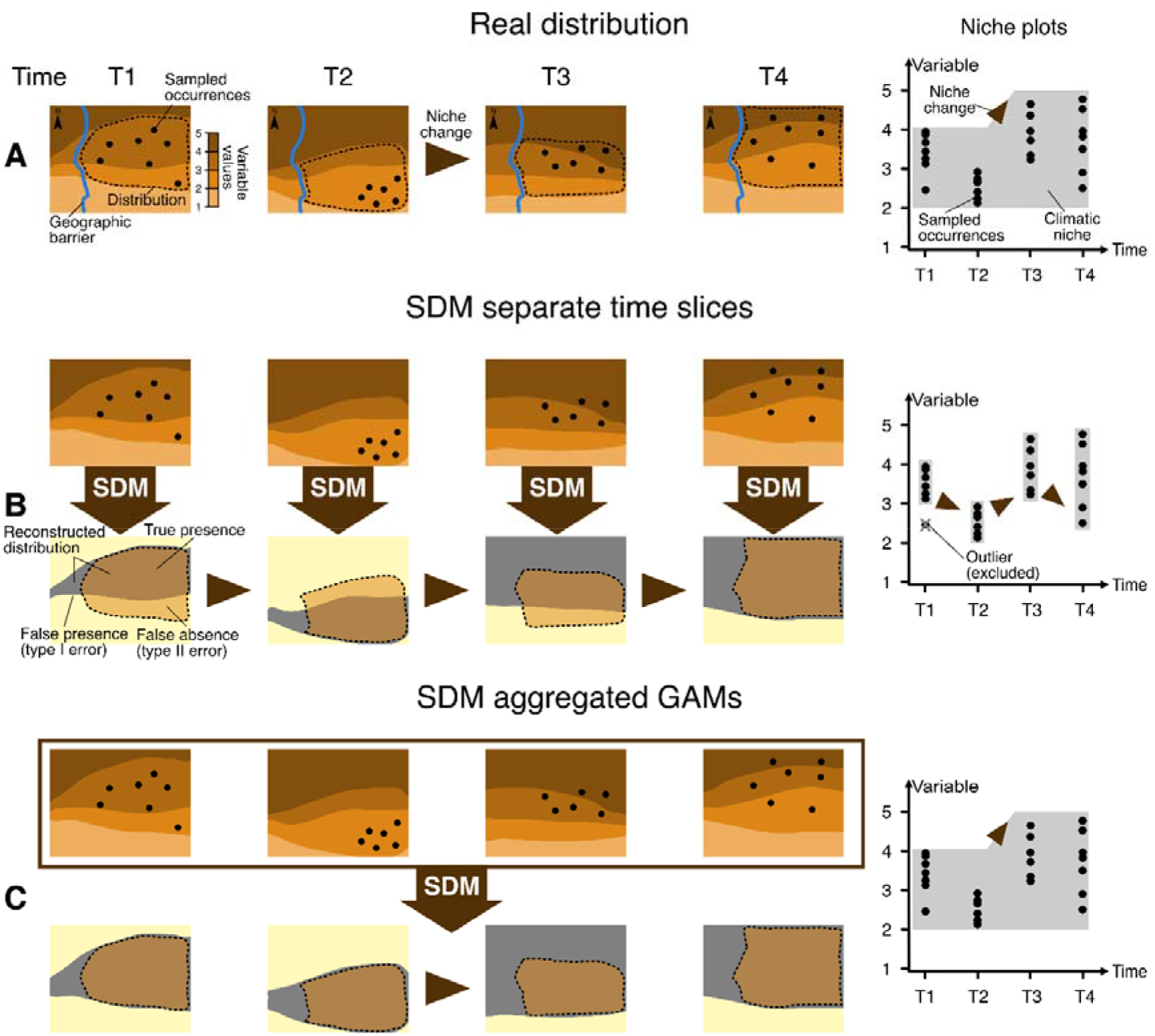
schematic description of different Species Distribution Modeling (SDM) approaches deal with diachronic data.

Human activities have strongly modified both the natural habitats (e.g. habitat loss, degradation, fragmentation), and the distribution of species (e.g. local extirpation). For this reason, the modern-day occurrences of a species may only represent a portion of their realized niche in terms of climatic variables, as many potentially viable areas may be practically unavailable due to anthropic pressures (2). A solution to this problem is a palaeobiological perspective that uses occurrences recovered from the archeological or paleontological record over thousands of years (2, 5). This approach helps to overcome the problem by defining the realized niche in the absence of the current anthropogenic pressures which, in turn, allows envisaging better conservation and restoration strategies (2, 5).

The last 50 thousand years (ky) provide an ideal time span to reconstruct the realized niche under different climatic conditions. Over this period, Europe has experienced very large climatic fluctuations (6) as it encompasses both the Last Glacial Maximum, when ice covered almost half of the continent, and the Holocene climatic amelioration that led to a drastic change of the vegetation composition of the continent (7). The last 50 ky also present a major technological advantage: they are covered by the radiocarbon dating method (^14^C, (8)), allowing to gather observations from the archaeological record with precise chronological attribution and associate them with palaeoclimatic reconstructions. The recent development of nearly-continuous palaeoclimatic data series over the last tens of thousands of years (9–11) provides the appropriate context to investigate species responses to climatic change over this time scales.

When SDMs have been applied to palaeoecological databases to reconstruct the niche through time, they have often been fitted independently for each time slice with enough occurrences and for which the paleoclimate was available (e.g. (12–16), and the niches estimated for each time slice compared to each other (13)(Figure 1B). However, this approach can be problematic, since the number of occurrences available for any time slice is often limited, compounded by a tendency for the sampling of the paleontological/archaeological record to be scarce, and often geographically incomplete (see the discrepancies between the sampled points and the outlined distribution in Figure 1A). Figure 1B illustrates how applying SDMs to each time slice independently may lead to substantial errors in the result, as a geographic bias in the samples may lead to undersampling the niche space in each time slice (Figure 1B, niche plots).

This undersampling, in turn, can have two important consequences. The first is an underestimate of the potential distribution of the species over space in each time slice, due to unsampled realized niche space because of sample bias. In order to correct this, it is important to aggregate the observations from different time slices before performing the analyses (17). The second consequence is that analyzing each time slice separately with sparse sampling may lead to an overestimate of the differences between them, identifying niche changes even when they have not occurred (Figure 1B). This is a particularly tricky aspect as there has been a long debate over niche plasticity within the time frame intermediate between geological and historical times (i.e. thousands of years) (18), which is yet to be resolved. On a deep time frame, niche evolution has been mainly analyzed in the light of diversification and speciation (19). At the other extreme, studies on the response to the ongoing climate emergency have identified several mechanisms allowing for plasticity in a very short time frame, such as changes in behavior (20, 21), phenology (22), epigenetics(23), but it is doubtful if this plasticity leads to long-term changes in the niche (24).

Traditionally, SDM have been developed for contemporary ecological data, so they have not been designed to explicitly take into account niche variation through time. In this work we propose to do so using the Generalized Additive Model (GAM) framework to analyze all available data within a single model. This method is already commonly used in SDM for static niches, but we adapt it to look at diachronic data allowing to test for changes in niche through time. This can be achieved by fitting interactions (technically tensor products) between predicting environmental variables and time, thus allowing the effect of those variables to change through time (Figure 1C). This approach alleviates the issue of patchy and limited sampling, as it can use the full time series to test for changes in the use of a certain part of the environmental parameter space (Figure 1C). In other words, we can take into account whether a species was present before and after a certain time point in that niche space, and use that information to avoid forcing a niche change if there is only a temporary absence due to limited data. This approach is already used in animal movement analysis to test for changes in habitat preferences (25, 26), an analogous question to that of niche change but focusing on a shorter time scale.

Here, we use this approach on four European megafauna species: wild horses (*Equus ferus*) and aurochs (*Bos primigenius*), which inhabited more open habitats, and wild boar (*Sus scrofa*) and red deer (*Cervus elaphus*, from now on *“deer”*), which are forest dwellers. All of them survived the Pleistocene-Holocene transition, which makes them ideal candidates to test whether their niche changed through time, and relate the timing of those changes to possible drivers. Furthermore, these species provide a range of representation in the archaeological record, and thus sampling completeness, with horse and deer being very common and aurochs and wild boar much less frequent.

## Results

### Occurrences from the paleontological and archaeological record

For horses, we used as occurrences an updated version of the radiocarbon dataset published in (12), removing points Eastwards of the Carpathians to only sample the European niche (12). For the other species, we collected from the literature and online databases direct or indirect radiocarbon dates (see Methods for details). We removed dates older than 40 thousand of calibrated years before present (kya), as there were too few data points for reliable analysis; to avoid sampling potentially domesticated occurrences, we also excluded occurrences younger than 8 kya (Figure 2).

**Figure 2:**
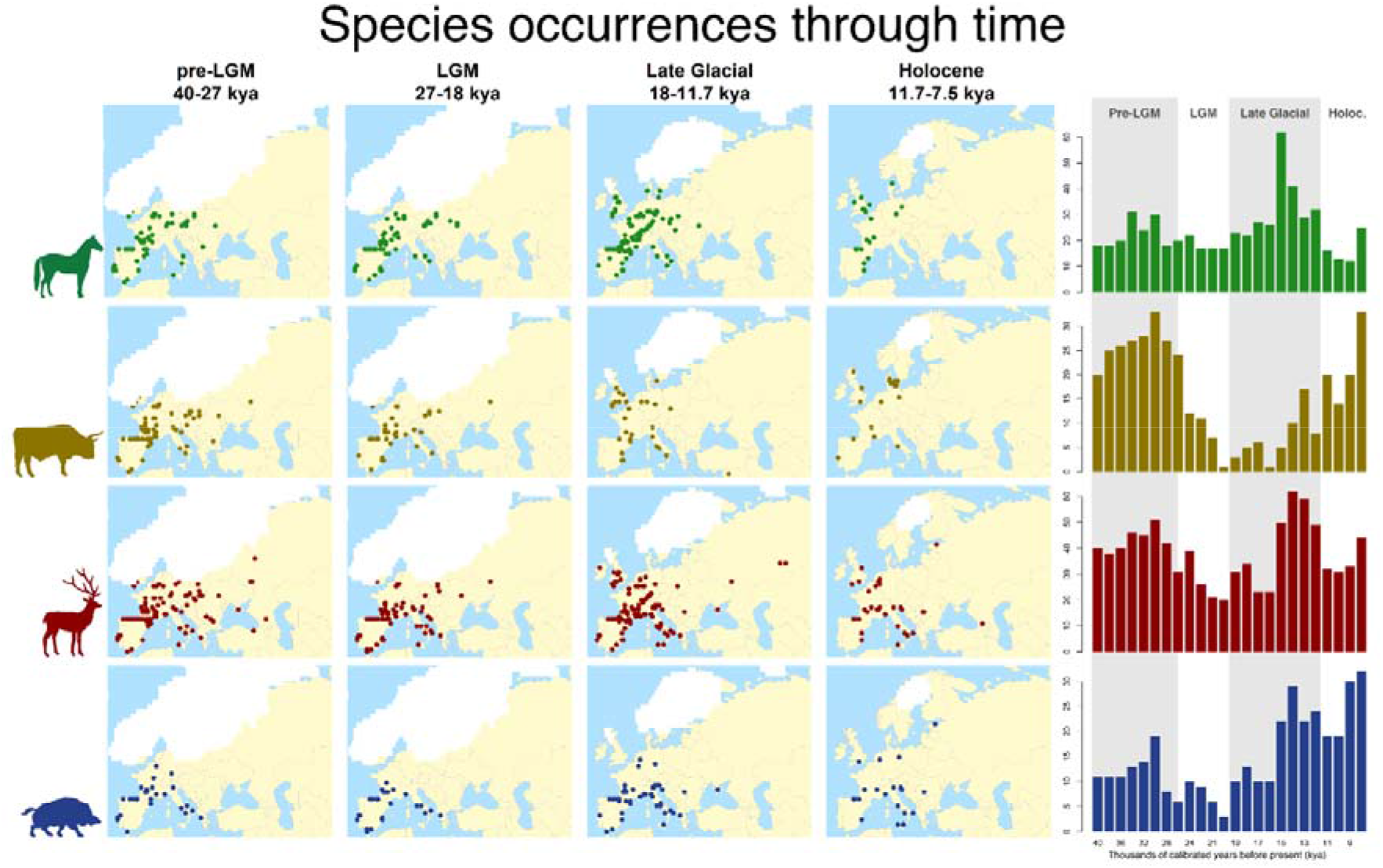
Geographic and temporal distribution of the occurrences analyzed respectively for horse, aurochs, deer and wild boar. Dates are expressed in thousands of calibrated years before present (kya).

For each occurrence point, we used its spatial coordinates and calibrated radiocarbon date to associate it with relevant palaeoclimatic variables (27). Palaeoclimatic reconstructions are available at a resolution of 0.5° for time slices of 1,000 year up to 21 kya and 2,000 years before that date. We focused on four variables which are relevant to large herbivores in Europe (see also (12)): maximum temperature of the warmest month (BIO5, from now on “maximum temperature”); minimum temperature of the coldest month (BIO6, from now on “minimum temperature”); total annual precipitation (BIO12, from now on “precipitation”) and net primary productivity (NPP). We performed a Principal Component Analysis (Figure S1) to visualize the niche space covered by the occurrences from each species over the whole time period, compared to the climate in Europe (baseline). Whilst there is a high degree of overlap in the distribution of the points in the four taxa, which is unsurprising given that they are all temperate herbivores, each species revealed subtle differences in their utilized niche, with deer showing the broadest one (Figure S1).

### Testing for niche change through time

We fitted two models to each species, a simple GAM using only the bioclimatic variables (“constant niche model”) and a full model including the interactions with time (“changing niche model”). GAMs require both absences and presences; so, we generated 20 sets of pseudo-absences that could be paired with the presences (i.e. our observed occurrences). In each set, for each presence, we generated 50 randomly located pseudo-absences with the same age (see Methods for details); thus, each model was fitted to 20 datasets of presences and different sets of pseudo-absences. We assessed the fit of the two models to the 20 datasets through calibration (28) using the Boyce Continuous Index (BCI) (29, 30), using an acceptance threshold of 0.8. Models for which at least 15 out of 20 repetitions passed such threshold (i.e. are robust to the random sampling of pseudoabsences) were then averaged in two different ensembles (by mean and median), and such ensembles were compared to see if the inclusion of an interaction with time in the analysis improved the fit to the data.

For horses and aurochs, the constant niche model failed to explain the data (i.e. fewer than 15 sets had a BCI >0.8; Table 1, Figures S2-S9), while for deer and wild boar it was able to describe the data adequately. Importantly, adding an interaction with time did not lead to an improvement of the fit for the latter two species, but it did generate models that could explain the distribution of horses and aurochs. We investigated the niche changes in these two species by plotting the averaged smooths from the GAMs with interactions (the individual ones are in Figures S10-S11), and contrasted this habitat suitability measure (Figure 3A) with the distribution of available habitat and presences over time (Figure 3B).

**Figure 3:**
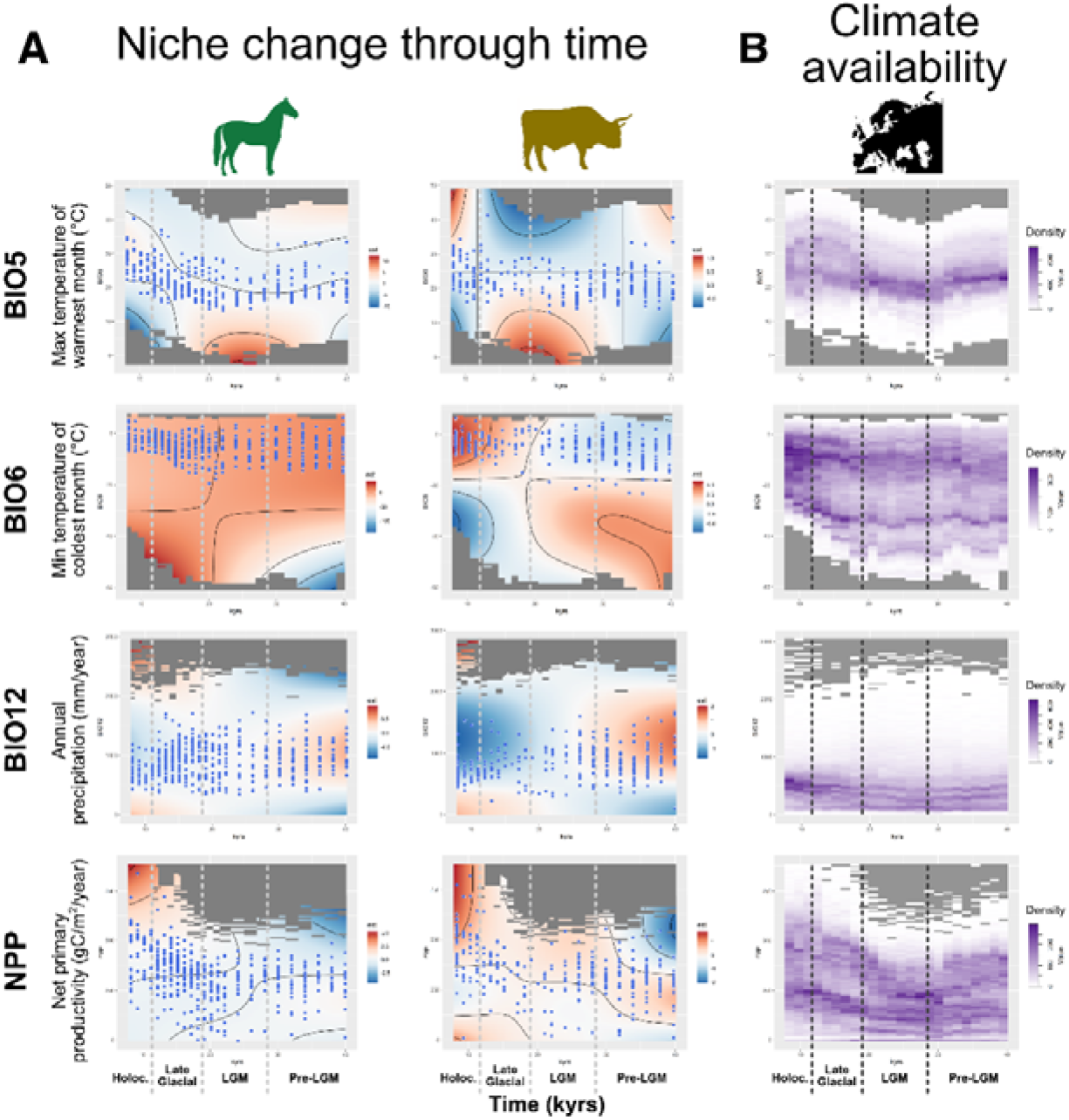
A: Depiction of how the niche changed through time through plots of the interaction between the climatic variables (y axis) and time (x axis). Red represent preference, and blue dislike. The points over the plots show the distribution of the species observations for each time slice. B: Climate availability, *i.e*. density of each variable within Europe through time (the darker the shade the higher the density). In all cases the variables are on the y axis and time on the x axis; the horizontal scales are consistent within each variable.

**Table 1.** Values of Boyce Continuous Index for the ensembles. Values in bold identify the best result for each species, which are the models used to predict the habitat suitability. N.A. represent models with less than fifteen pseudoabsence runs above the acceptance threshold, for which the ensemble has not been built.

| Models | Horse |  | Aurochs |  | Deer |  | Wild boar |  |
| --- | --- | --- | --- | --- | --- | --- | --- | --- |
|  | mean | median | mean | median | mean | median | mean | median |
| Constant niche (simple model) | N.A. | N.A. | N.A. | N.A. | **0.98** | 0.97 | **0.90** | 0.65 |
| Changing niche (full model) | **0.92** | 0.79 | **0.94** | 0.89 | 0.93 | 0.75 | N.A. | N.A. |

For BIO5, BIO6 and NPP, both horse and aurochs tended to track the most available values (dark purple in Figure 3B). This means that for maximum temperature (BIO5), their distribution mostly spanned between 15° and 35° C, with a tendency towards lower values during the LGM and higher values during both the pre-LGM and the Holocene. Minimum temperature (BIO6) showed a bimodal distribution of its availability in Europe, following through time two main trajectories of high frequency values (Figure 3B). Both species tended to follow the upper trajectory (approximately between 10° and −20°C). The aurochs tended to occupy a smaller variable space than the horse, with its range shrinking more during the Holocene. Precipitation (BIO12) is the only variable for which our species did not occupy the part of the range which is most available in the space (below and around 500 mm per year), inhabiting areas between 500 and 1500 mm per year. There was also a range reduction in the most recent period, especially for horses. For NPP, horses track the two high-frequency trajectories, while aurochs mainly follows the upper one. This means that, in both species, the occupied variable range includes lower values during the LGM and then higher ones in the following periods.

### Potential distribution through time

We used the best ensemble for each species to project the potential distribution over each time slice, and generated binary maps using the highest threshold allowing to recover 99% of our data (Figures S12-15). We then averaged the binary distribution for the individual time slices over each climatic period (Figure 4). The potential distribution of the four species was generally restricted to Central, Western and Southern Europe. During the pre-LGM, horse and wild boar occupied only Central and Southern Europe, while the range covered by aurochs and deer extended more towards East, especially in the latter species. During the LGM, the ranges contracted for all species but deer, which showed the same potential distribution between the pre-LGM and the Late Glacial. Starting from the Late Glacial the pattern became more similar for horses, aurochs and wild boar, with an expansion first towards north, and then east during the Holocene. On the other hand, the distribution of deer only expanded during the latter period, reaching far more eastern regions than the other species.

**Figure 4:**
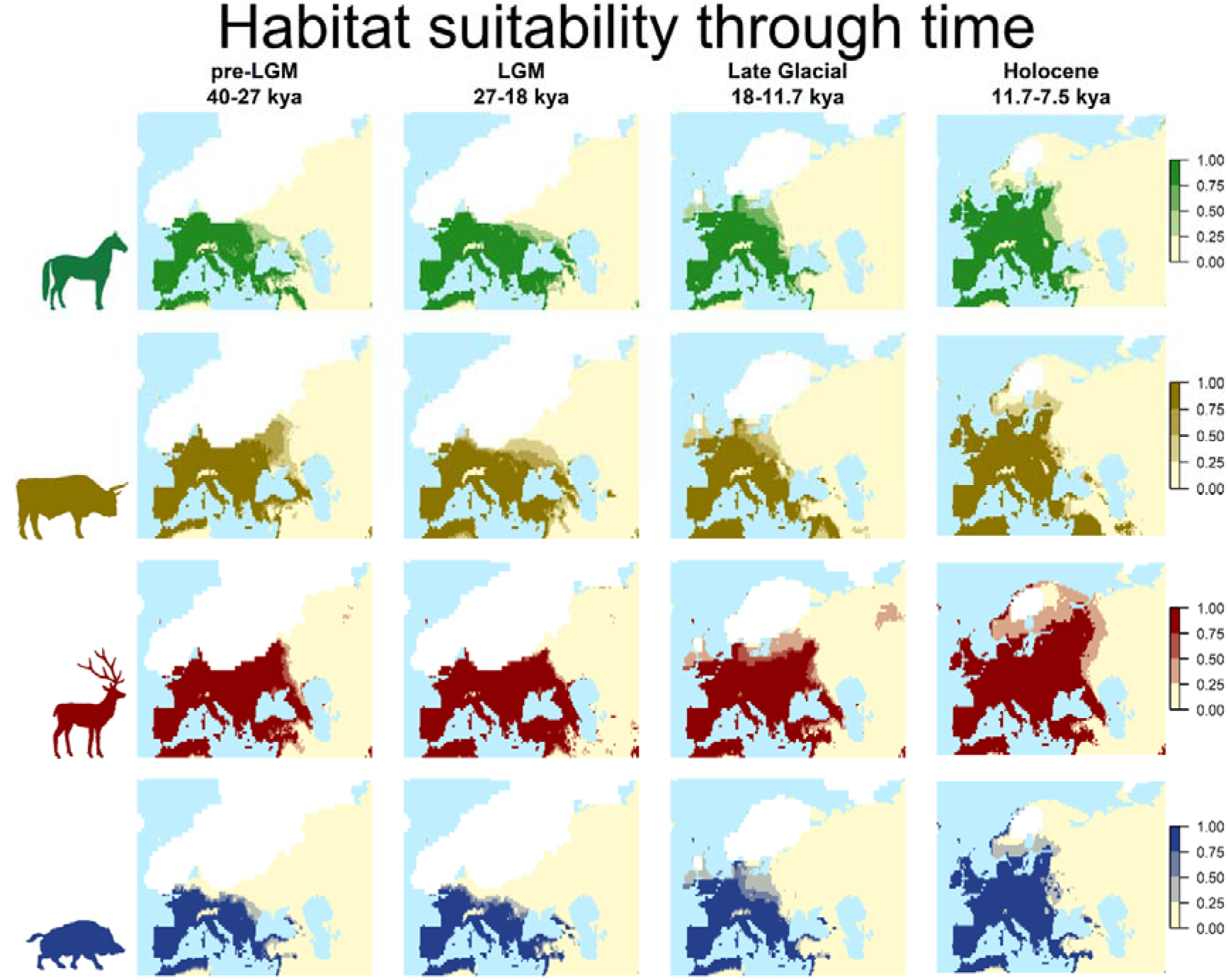
Potential distribution of the species for each climatic period. They are created averaging the distributions for all time slices within each period, calculated using the best ensemble for each species.

### Niche analyses

We formally quantified niche overlap between pairs of species by calculating Schoener’s D (31, 32), merging the reconstructed binary distributions from each time slice over the whole time-frame considered (Table 2). Horse and aurochs, both specialists of open habitats, showed the highest overlap, and they differed greatly from deer, which is a forest specialist. Wild boar, which is able to use semi-closed habitats, appeared to be intermediate among all species in its niche requirements.

**Table 2:** Niche overlap (Schoener’s D) between species. Darker filling is associated with higher values

|  | Horse | Aurochs | Deer |
| --- | --- | --- | --- |
| Aurochs | 0.90 |  |  |
| Deer | 0.73 | 0.73 |  |
| Wild Boar | 0.81 | 0.82 | 0.80 |

We also analyzed how climatic changes influenced the realized niche in each time slice by calculating Schoener’s D between adjacent time slices for each species. As shown in Figure 5A, the level of overlap through time varied between 0.5 and 0.9, with values between 0.5 and 0.7 considered “medium”, and above 0.7 “high” (33). The pattern is very similar among species, with lowest values observed at the transitions between 34 and 32 kya, and 12 and 11 kya. Deer distinguish themselves from the others by showing lower values also between 28 and 22 kya, and between 16 and 14 kya.

**Figure 5:**
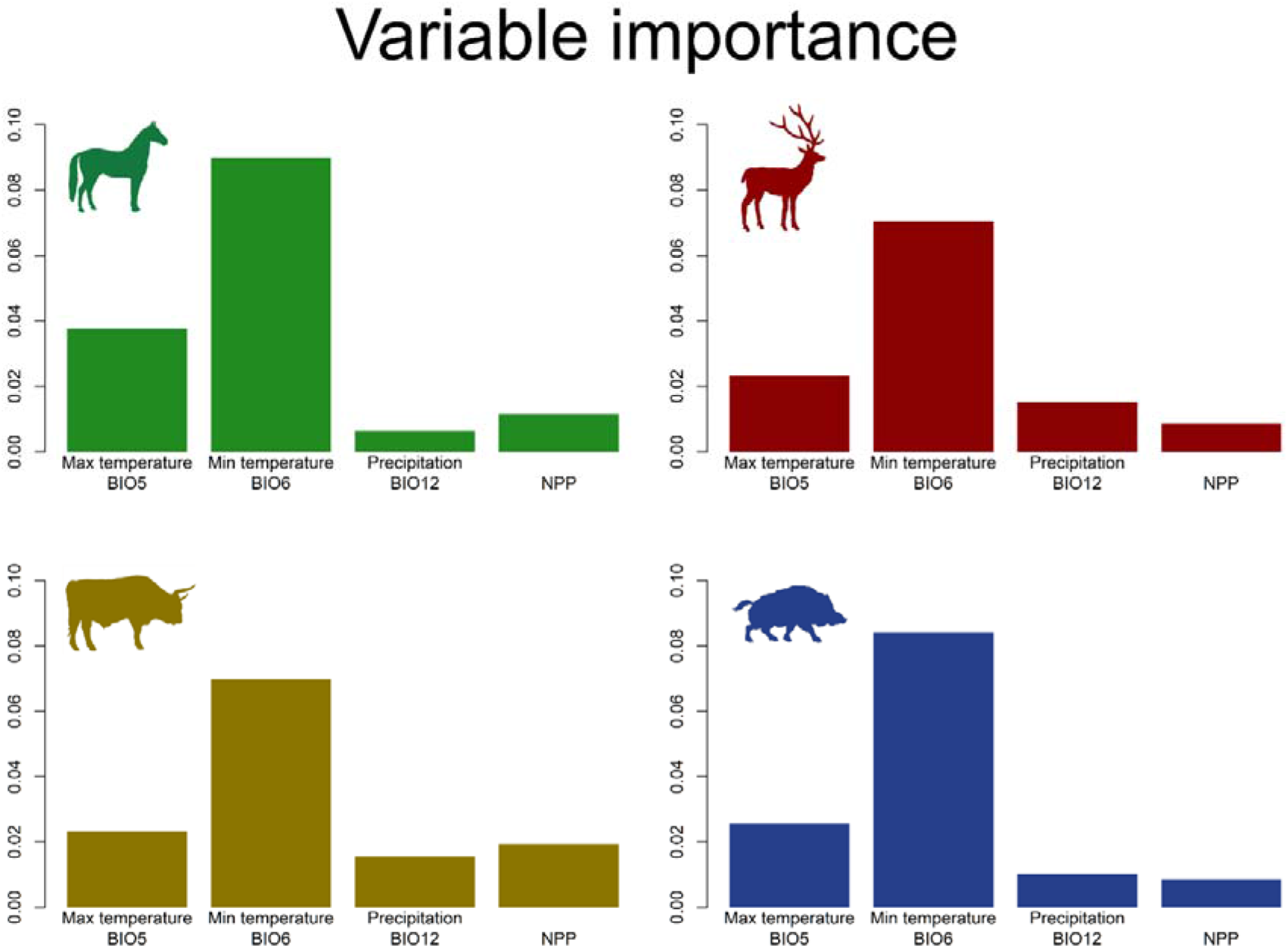
Variable importance.

Finally, we calculated the variable importance (Figure 5), represented by the unique deviance explained by each of them. The pattern is consistent in all species, with minimum temperature (BIO6) showing the highest value, followed by maximum temperature (BIO5). Among the two remaining variables, NPP is more important than precipitation (BIO12) for horse and aurochs, while the opposite is true for deer and wild boar.

## Discussion

The reconstructed niches for the four species showed levels of similarity in accordance to their ecology. The niches of horse and aurochs overlapped to a large extent, whilst being distinct from that of deer. Wild boar had an intermediate niche, likely associated with its high degree of omnivory compared to the grazing and browsing requirements of the other three ungulates. Furthermore, NPP played a greater role for the for the open habitat species, whilst precipitation was more important for the close habitat ones. Despite these differences, all four species were predicted to occupy large parts of Western Eurasia, showing a degree of overlap both geographically and in niche space. Their broad climatic niches likely played a role in their success as “winners” that survived the megafauna extinctions that occurred during and immediately after the Last Glacial Maximum.

Despite the patchy and limited sampling of archeozoological remains, we were able to capture the differential role of mountains as barriers for these species. For the open habitat species, horse and aurochs, the Alps and Caucasus were unsuitable throughout the study period. This reconstruction is consistent with mountains acting as genetic boundaries for this species as it has been suggested with the Alps for aurochs (34, 35) and with the Pyrenees for horses (12, 36, 37). For the other two species, on the other hand, these major mountainous areas were only unsuitable during the colder periods, but became suitable habitat with the warming of the Holocene, in line with the expansion of forests in these areas. We would caution against overinterpreting the lack of signal from other mountainous regions; on one hand, it might derive from their lower altitudes (e.g. the Carpathians or the southern Balkans), but the inevitably coarse scale of our reconstructions might have also prevented us from detecting mountains with a smaller footprint such as the Pyrenees.

Compared to the other species included in this study, horses are better adapted to colder environments and are considered a typical element of the steppe-tundra community (38, 39). This observation might, at first, seem discordant with our results, as their reconstructed distribution is highly overlapping with aurochs and, to some extent, wild boar, two species which are more characteristic of temperate environments (39). This lack of an expansion towards colder regions is in fact the result of our focus on Europe. Previous SDM analyses on horses have showed that this species occupied two separate niches: a colder one associated to Asia and part of Eastern Europe, and a warmer one covering instead Central, Western and Southern Europe (12). By limiting our observations to the West of the Carpathian Mountains (Supplementary figure 16), we have focused on the warmer niche. When part of the eastern range was included (Supplementary figure 17), the potential distribution of the species stretched towards North-East, partially recovering the Asian range. For Europe, the archaeological record from Holocene sites (12) suggests that, even though the species might have at time coexisted, horses were relatively rare when the other two taxa were abundant, highlighting that an overlap in range does not necessarily imply identical habitat preferences.

Deer has a wider potential distribution compared to the other species, stretching more towards east for the whole period analyzed. This is consistent with previous archaeological analyses: compared to aurochs and wild boar, deer has been more tolerant to different habitats, and in warmer periods it co-occurred with species from the steppe-tundra biome (e.g. mammoth, reindeer, cave bear) (39–41). Interestingly, the reconstructed range did not shrink during the LGM, predicting the presence of this species further north than the southern areas that have been suggested by both the archaeological record (42) and genetic analyses (43). An analogous pattern, albeit less pronounced, is found for the other three species. Their predicted ranges during the LGM encompass what are considered to be glacial refugia for temperate species during this period (44–46): the Iberian, Italian and Balkan Peninsulas, Dordogne, and the Carpathians. However, our reconstructions cover a slightly larger area extending to regions that are not known to be inhabited by the analyzed species (e.g. most of Germany and the Carpathians for wild boar) (44).

There are several possible explanations for this discrepancy. First of all, when looking at northern France and Germany, there is very limited archaeological evidence for the LGM (47). Thus, a lack of observations might be a false negative due to inadequate sampling effort. The mismatch between predictions and observations could also be linked to such marginal regions being ecotones, i.e. transitional areas between two ecologically different zones. From a climatic point of view, they could be easily occupied by our species, but both the animal and vegetal communities would be a blend of the two neighboring ones. This would cause biological interactions (e.g. competition, predation) that could lead the species being less frequent than in their core areas, and thus less observed in the archaeological record. This effect could be further compounded by differential predatory choices by human populations through space and time, a factor that may heavily influence the presence or absence of species in the archaeological assemblages. For instance, our previous SDM analyses on horse (12) showed that, although during the Holocene a large part of Europe was still climatically suitable for the species (and it was apparently still widespread throughout the continent), this species is mostly absent from the zooarchaeological record and its presence in faunal assemblages is usually less than 3 % of the identified remains. A further complication is that our climatic reconstructions are inevitably coarse (each cell is approximately 100km wide); whilst the average climatic conditions might be suitable for the species, heterogeneities in microclimate might lead to habitat fragmentation that could greatly reduce species density, or even preclude the viability of populations. A similar issue was discussed in (12, 48): when horse remains were found in areas where the macroclimate suggested forested environment, they were shown to be living in open areas based on microhabitat reconstructions based on the faunal assemblages at archaeological sites.

Our analysis only considers climatic factors, and both dispersals and biotic interactions are not evaluated. Beside potentially missing certain changes in niche (51), it is also important to note that observed changes in the bioclimatic niche could be indirect effects of some of these unmeasured factors. Indeed, this might be the case for the change in niche for our two open-habitat species (horse and aurochs), which we detected at the transition from the Last Glacial Maximum to the Holocene (whilst no change is found for the two forest species, deer and wild boar). The majority of megafauna extinctions following the LGM affected large open habitat herbivores and their predators, potentially reshaping the communities in this habitat. (49, 50); thus, the observed change in niche might be the result of the removal of competitors and predators. Moreover, in the same time frame human populations changed their hunting behavior (52), potentially affecting differently various animal species.

By estimating the realized niche of the species over a long period of time with large climatic fluctuations, our approach has multiple strength. By using the archeozoological record, it overcomes the bias given by contemporary occurrences that may be strongly influenced by human pressures (2). At the same time, archaeological and radiocarbon datasets suffer of important geographic biases because historically some regions (e.g. Central Europe) have been subject to more extensive excavation and dating efforts than others (e.g. Eastern Europe). Our method, by taking into account all occurrences through time, is less sensitive to this problem, which is also reflected by the fact that the projected distributions include geographic areas that are strongly under-sampled in term of occurrences (e.g. Caucasus, Anatolia, etc.). Finally, and more importantly, our approach provides the best possible approximation of the fundamental niche of a species while testing for changes in the realized one.

From an evolutionary perspective, identifying changes in the realized niche is a key starting point for testing whether they are linked to specific adaptations (e.g. using ancient DNA data) or instead are shifts within a large fundamental niche. Both of these aspects could be highly significant when planning conservation or restoration efforts.

## Materials and methods

### Materials

For horses we used an updated version of the database published in (12), whilst for the other three species we collected direct and indirect radiocarbon dates from the literature. We only considered dates located in Europe (West of 60°E and North of 37°N).

All dates were calibrated with OxCal (8) version 4.3 using the IntCal20 curve (53) and then classified into time slices. We decided to only keep observations between the time slices of 40 and 8 kya: their scarcity before this period may bias the GAMs, and after it, domesticated cattle, pigs and (later) horses arrived to Europe making it difficult to differentiate them from their wild forms.

The datasets were cleaned by removing records without a reported standard error, and dates with calibration errors, resulting in the number of observations presented as “Raw dataset” in Table 3Table 3.

**Table 3.** Species datasets before (“whole”) and after (“final”) cleaning. The number reported for horses only includes the samples East of 28°E used for the analyses presented in the main text.

|  | Horse | Aurochs | Deer | Wild boar |
| --- | --- | --- | --- | --- |
| Raw dataset | 1088 | 757 | 1778 | 663 |
| Cleaned dataset | 575 | 396 | 933 | 368 |

For horses, we additionally removed points east of the Carpathian Mountains, identified with their easternmost limit of 28°E (Figure S16). European and Asian horses have been shown to have different niches (12). When we included dates east of the Carpathians, we recovered a potential distribution (Figure S17) that extended well beyond that reconstructed in (12), suggesting that we started to sample the Asian niche.

Data were then collapsed by keeping only one point per grid cell per time slice, leaving the number of observations reported in Table 3 as “Cleaned dataset”.

We used the paleoclimatic reconstructions published in (11), which are based on the Hadley CM3 model, choosing the same BIOCLIM variables as in (12): BIO5 (maximum temperature of the warmest month), BIO6 (minimum temperature of the coldest month), BIO12 (annual precipitation) and Net Primary Productivity (NPP), after checking for collinearity between them (correlation < 0.7 in all cases). Reconstructions are available for each millennium up to 21 kya, and every two millennia before that date, with a resolution of 0.5°.

The results of our analyses are presented by climatic periods, identified as follows: pre-LGM (from the beginning of the period analyzed, i.e. 40 kya, to 27 kya), LGM (from 27 to 18 kya), Late Glacial (from 18 to 11.7 kya), Holocene (from 11.7 kya to the end of the period analyzed, i.e. 8 kya).

### Methods

To visualize the niche of each species, we performed a PCA of the occurrences of the four species using the *princomp* function in R (*package stats*) against a reference set created by merging 5,000 points randomly selected from each time slice (and thus capturing the range of climates available through the time period of interest).

Given that absences from the sampled archaeological record are as geographically biased towards Central and Eastern Europe as the presences, and they do not cover the whole space available, we used pseudoabsences to adequately represent the existing climatic space in our SDMs. For each species, we generated 20 set of pseudoabsences. Each set was generated by sampling, for each observation, 50 random locations matched by time. This resulted in n=20 datasets of pseudoabsences and presences for each species, which we used to repeat our analyses to account for the stochastic choice of pseudobasences. For each dataset, we used GAMs to fit two possible models: a “changing niche” model that included interactions of each environmental variable with time (fitted as tensor products), and a “constant niche” model which included the environmental variables as covariates but lacked interactions with time. For all GAMs, we set 4 as maximum threshold for the degrees of freedom of the splines; this value provides a reasonable compromise between allowing the relationship to change through time but avoids excessive overfitting (25). GAMS were fitted using the *mgcv* package in R (54).

Model fit for each of the GAM was evaluated with the Boyce Continuous Index (BCI, 15, 16), which is specifically designed to be used with presence-only data and is well suited to perform model selection (28, 55), setting a threshold of Pearson’s correlation coefficient]>] 0.8 to define acceptable models (29). For each combination of species and model type (changing niche vs. constant niche), we assessed whether at least 75% of repeats (15 out of 20) passed the BCI>0.8 threshold. The relative importance of each environmental variable was quantified by computing the average of the variance explained by each variable over all the models above the BCI threshold of 0.8.

To achieve more robust predictions (56), we averaged in two different ensembles the repetitions above the acceptance threshold obtained for each model: by mean and median. This step is intended to reduce the weight of models that are highly sensitive to different pseudoabsence random samplings. For each species, we then selected the ensemble (either based on mean or median) with the higher BCI as the most supported, and use it to perform all further analyses.

To visualize the prediction for each species, we then transformed the predicted probabilities of occurrence from the ensemble into binary presence/absences by using the threshold needed to get a minimum predicted area encompassing 99% of our presences (function ecospat.mpa from the ecospat R package (57)). The binary predictions were then averaged over each of the major climatic periods.

The effect of different variables through time was visualized by plotting the interactions of the GAMs for the two species that showed a change in niche. For each model with a BIC>0.8, we used the R package gratia (58) to generate a surface with time and the environmental variable as the x and y axis, and the effect size as the z. For each species, we then plotted the mean surface, which captures the signal consistent across all randomized pseudoabsences.

To see how much the realized niches of the four species overlap, we aggregated in a single table the binary predictions from all time slices for each of them. We then used the R package ecospat to calculate Schoener’s D (31, 32) between each pair, following the standard procedure. The overlap is considered to be low between 0 and 0.3, medium between 0.3 and 0.7 and high between 0.7 and 1 (33).

## Acknowledgements

ML and AM have been funded by the ERC Consolidator Grant 647787 “LocalAdaptation”. The authors thank Elisa Anna Fano and Mario Coiro for their helpful comments on the manuscript and Robert Sommer for the useful discussion of the results.

## Author contributions

ML and AM designed the study with inputs from FB and PB. FB, PB and ML collected the data and curated the dataset. ML performed the analyses under the supervision of AM and designed the figures with inputs from all other authors. ML, FB and AM provided the interpretation with inputs from PB. ML and AM wrote the paper with inputs from all other authors.

## Notes

### Competing Interest Statement

The authors have declared no competing interest.

