## Supplementary figures for "Following The Niche: Reconstructing 32,000 Years Of Niche Dynamics In Four European Ungulate Species"

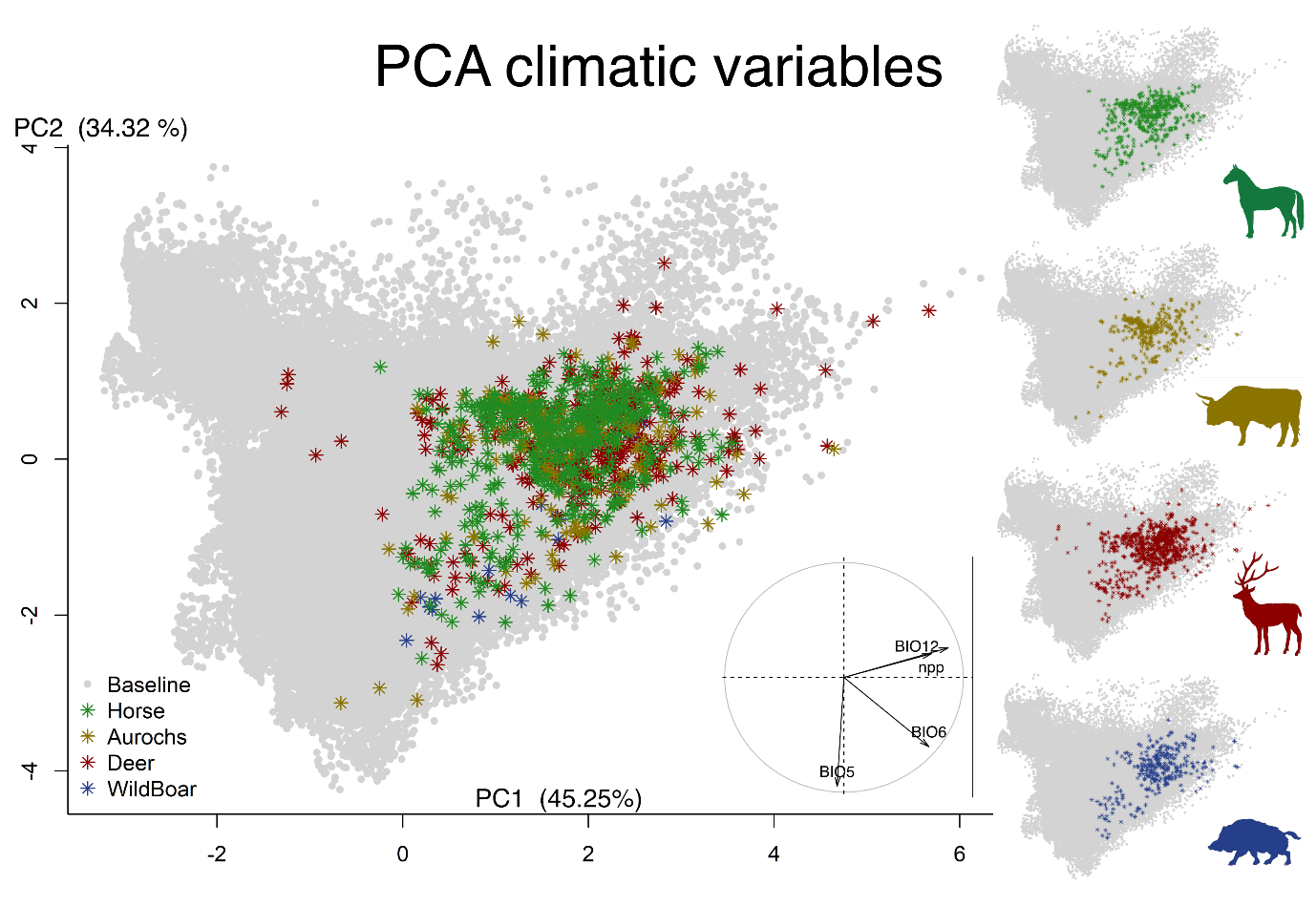

Figure S 1: Principal Component Analysis (PCA) of the four species occurrences (color asterisks) compared to the whole of Europe (Baseline, light gray dots).

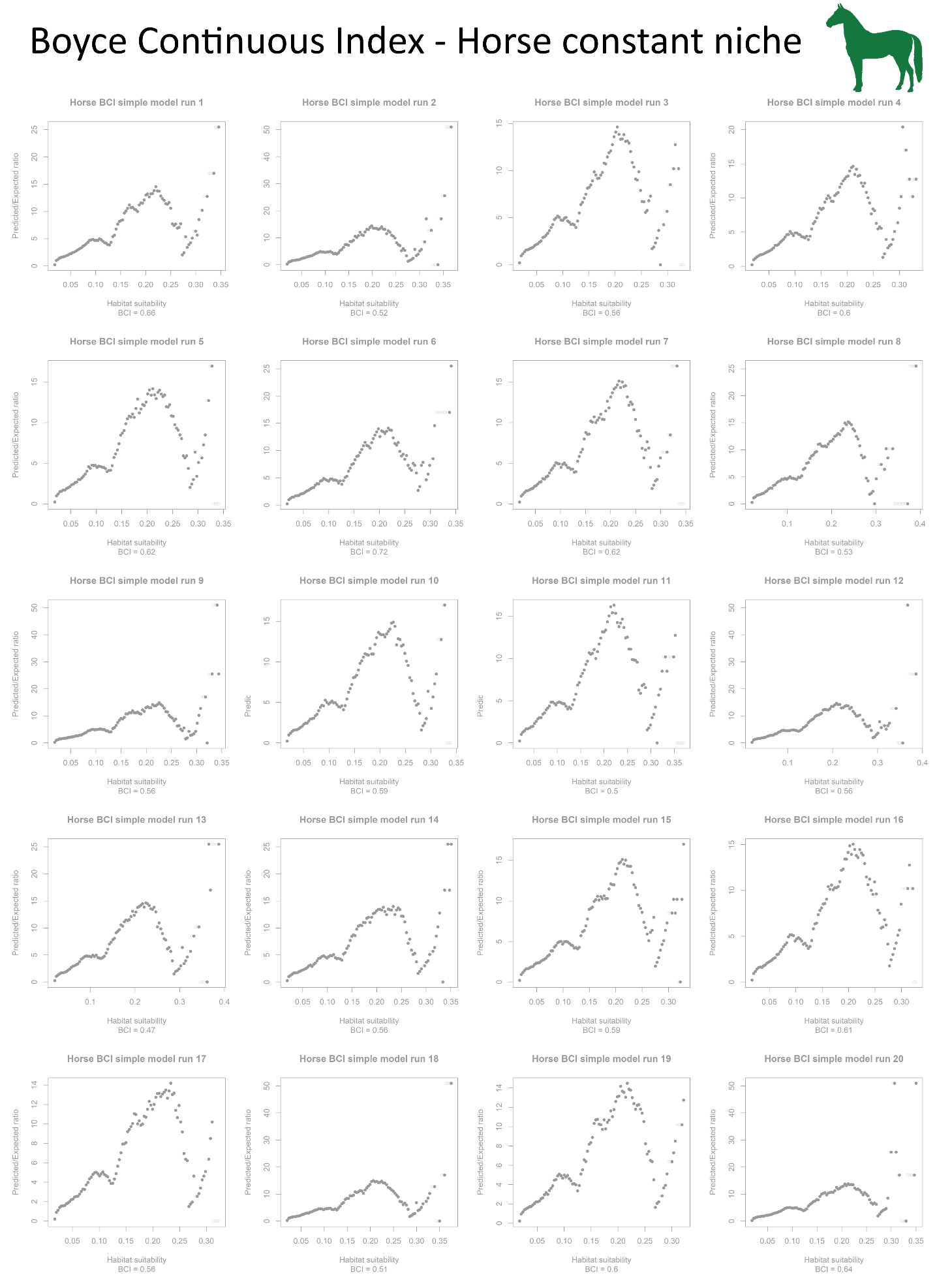

Figure S 2: Boyce Continuous Index (BCI) plots for each run of the constant niche model (= simple model) in horses. In gray, runs with a BCI below the acceptance threshold (0.8)

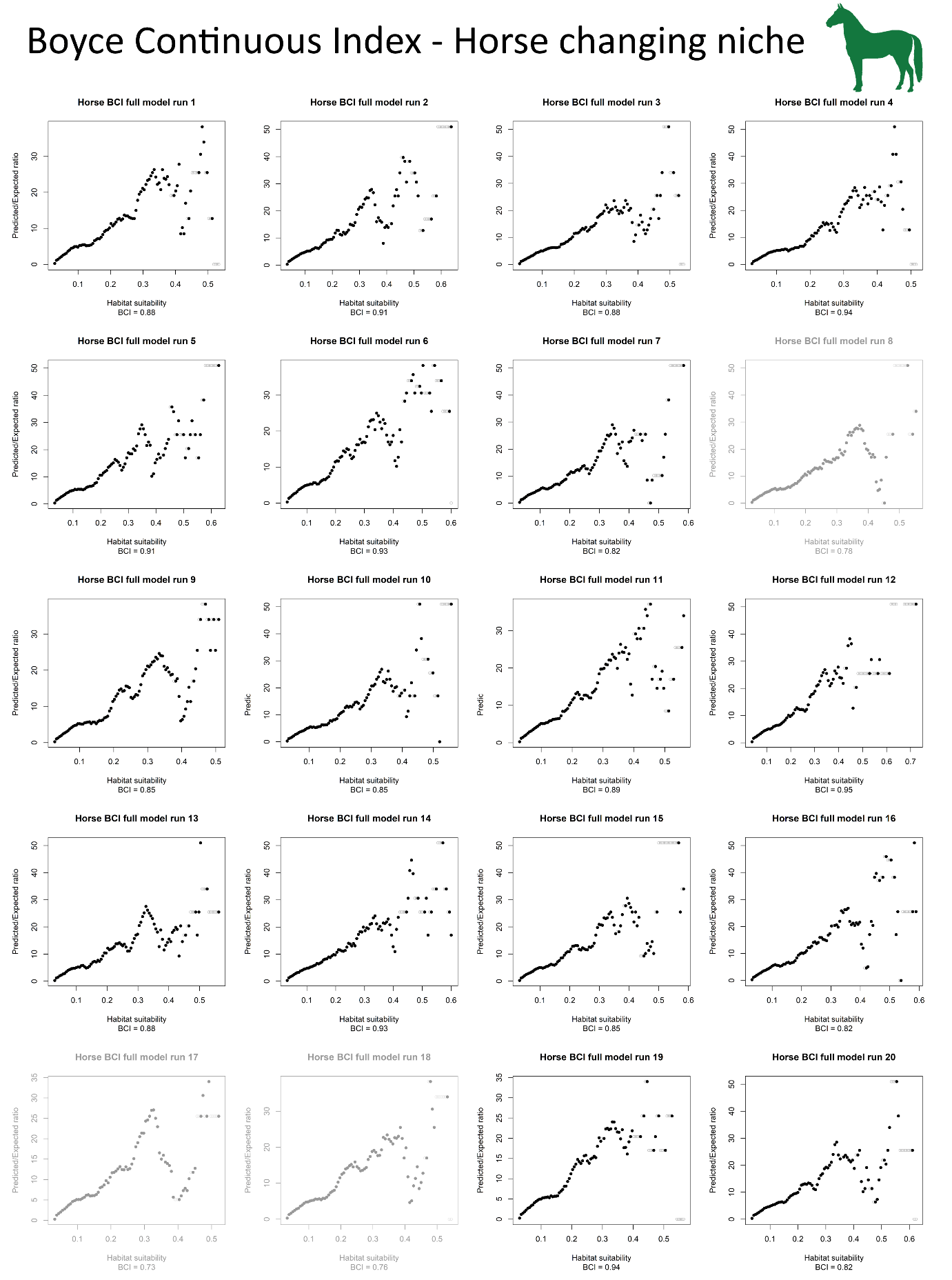

Figure S 3: Boyce Continuous Index (BCI) plots for each run of the changing niche model (= full model) in horses. In gray, runs with a BCI below the acceptance threshold (0.8).

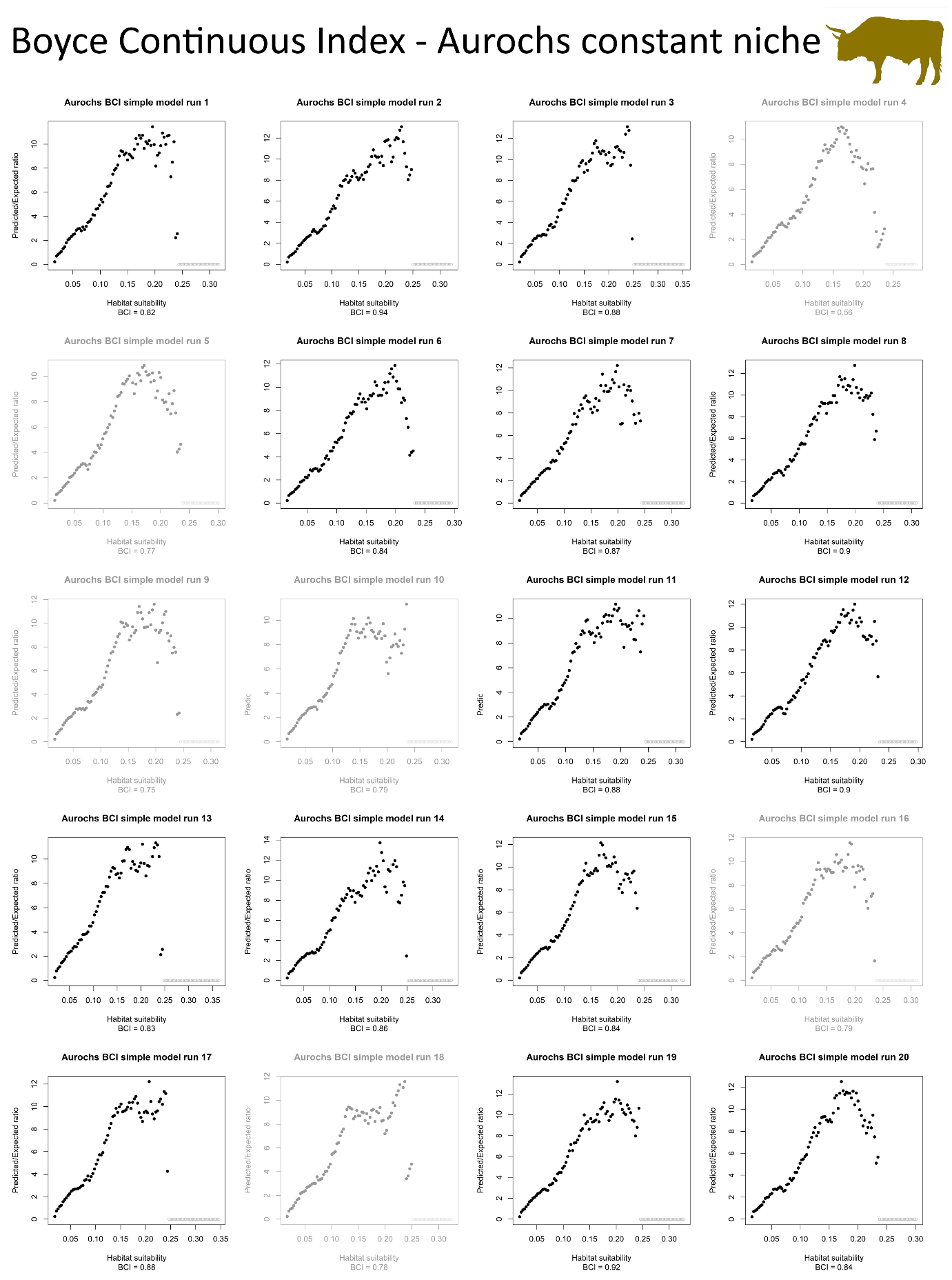

Figure S 4: Boyce Continuous Index (BCI) plots for each run of the constant niche model (= simple model) in aurochs. In gray, runs with a BCI below the acceptance threshold (0.8).

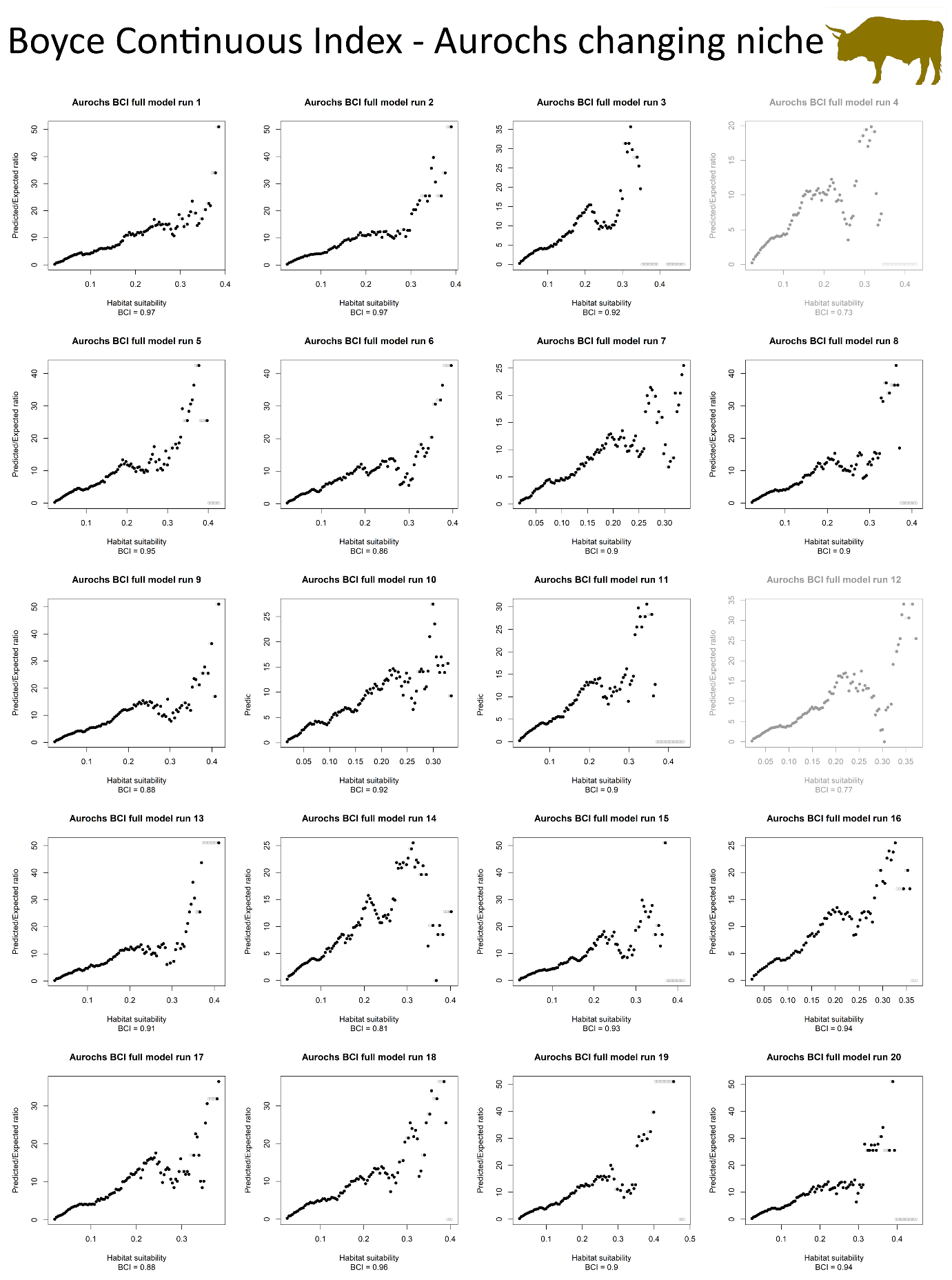

Figure S 5: Boyce Continuous Index (BCI) plots for each run of the changing niche model (= full model) in horses. In gray, runs with a BCI below the acceptance threshold (0.8)

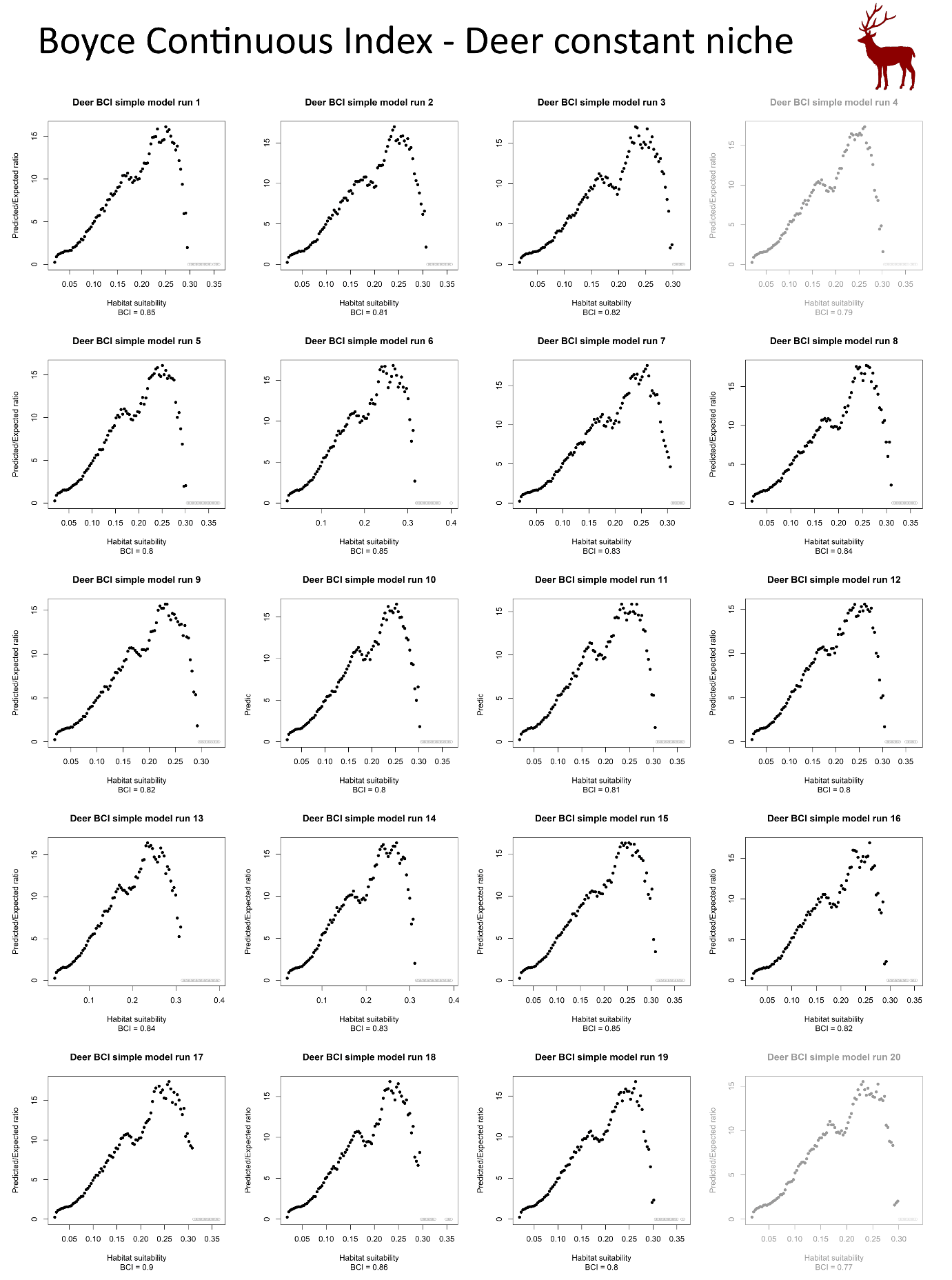

Figure S 6: Boyce Continuous Index (BCI) plots for each run of the constant niche model (= simple model) in deer. In gray, runs with a BCI below the acceptance threshold (0.8)

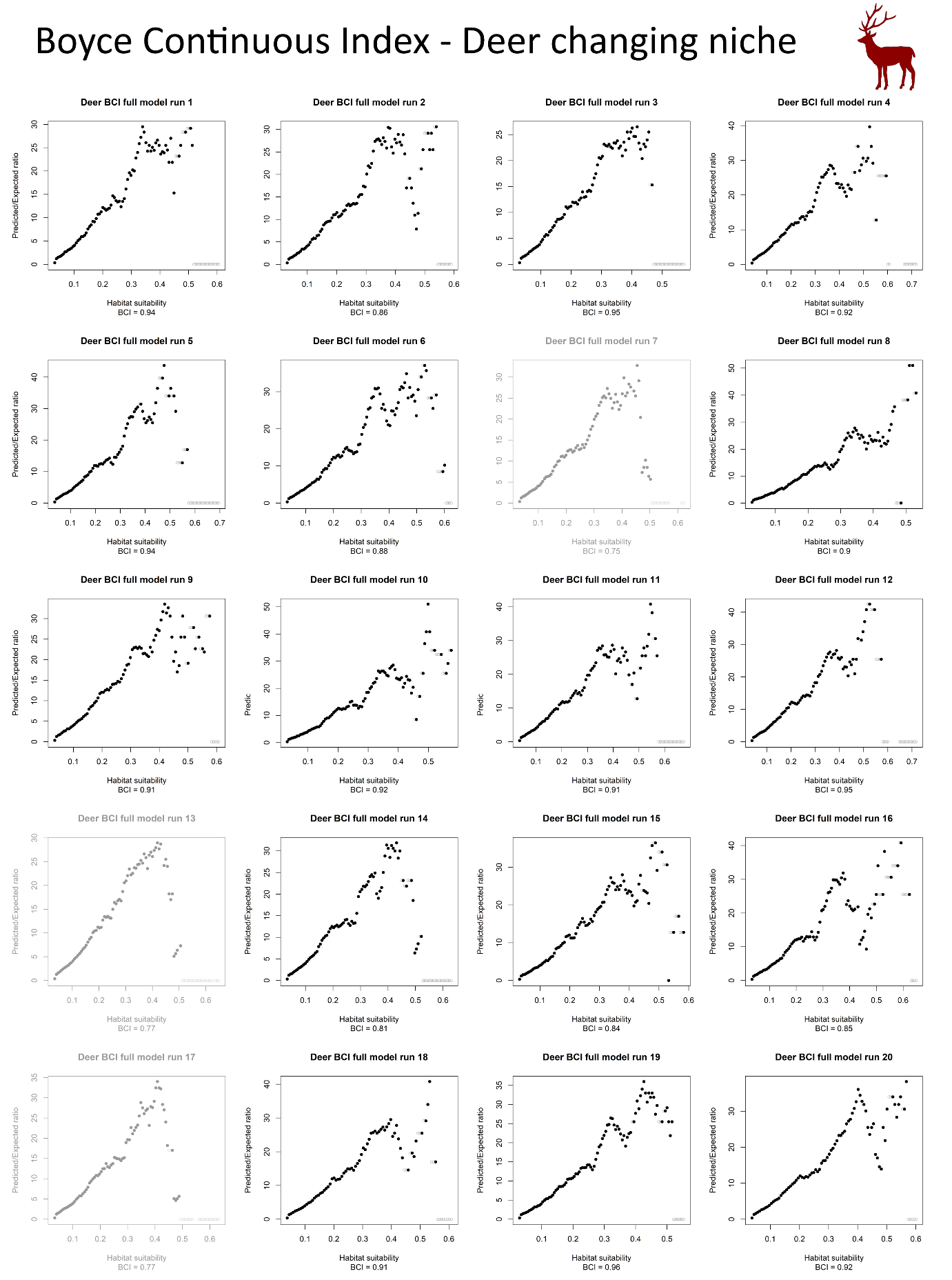

Figure S 7: Boyce Continuous Index (BCI) plots for each run of the changing niche model (= full model) in deer. In gray, runs with a BCI below the acceptance threshold (0.8)

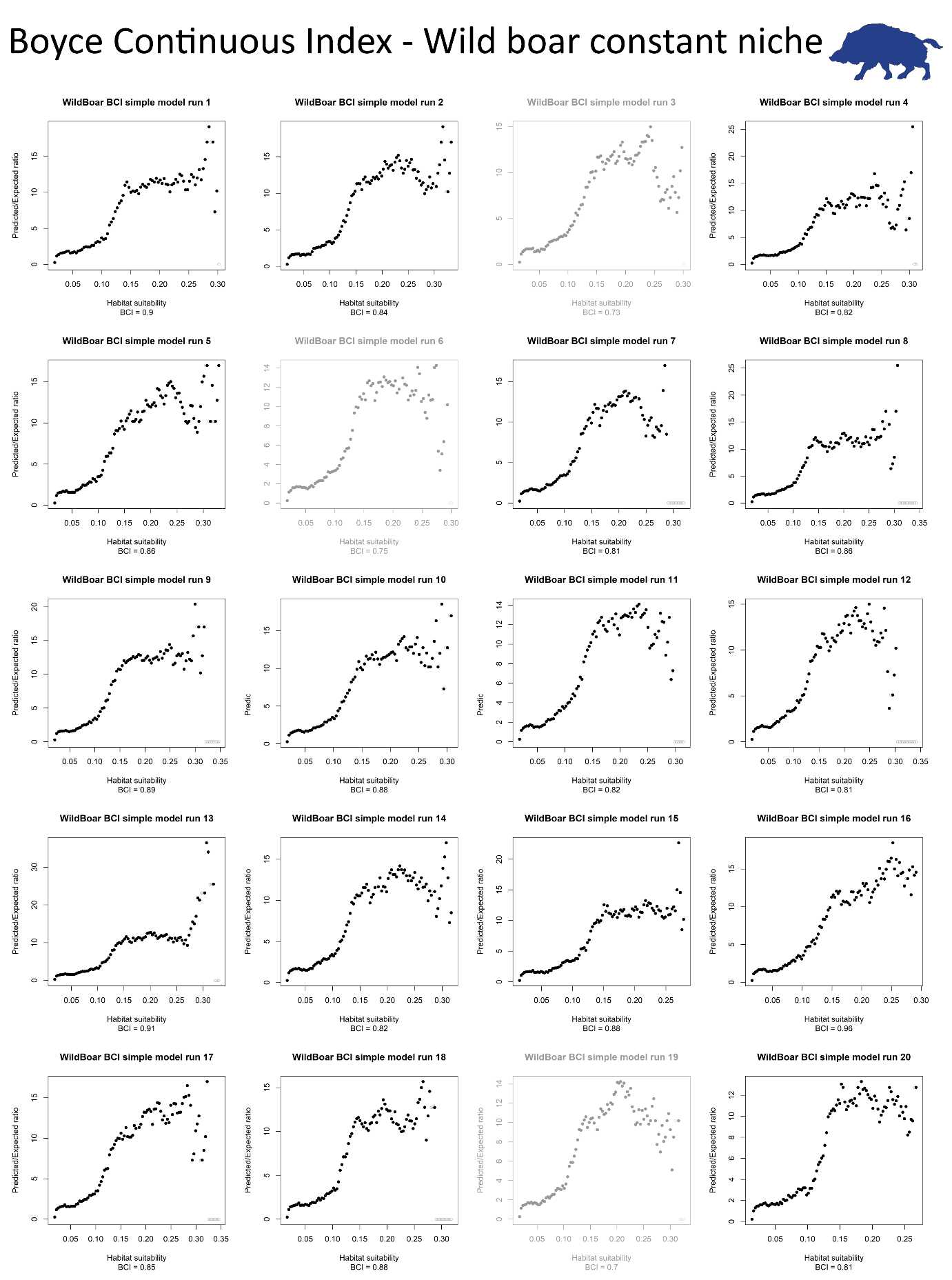

Figure S 8: Boyce Continuous Index (BCI) plots for each run of the constant niche model (= simple model) in wild boar. In gray, runs with a BCI below the acceptance threshold (0.8)

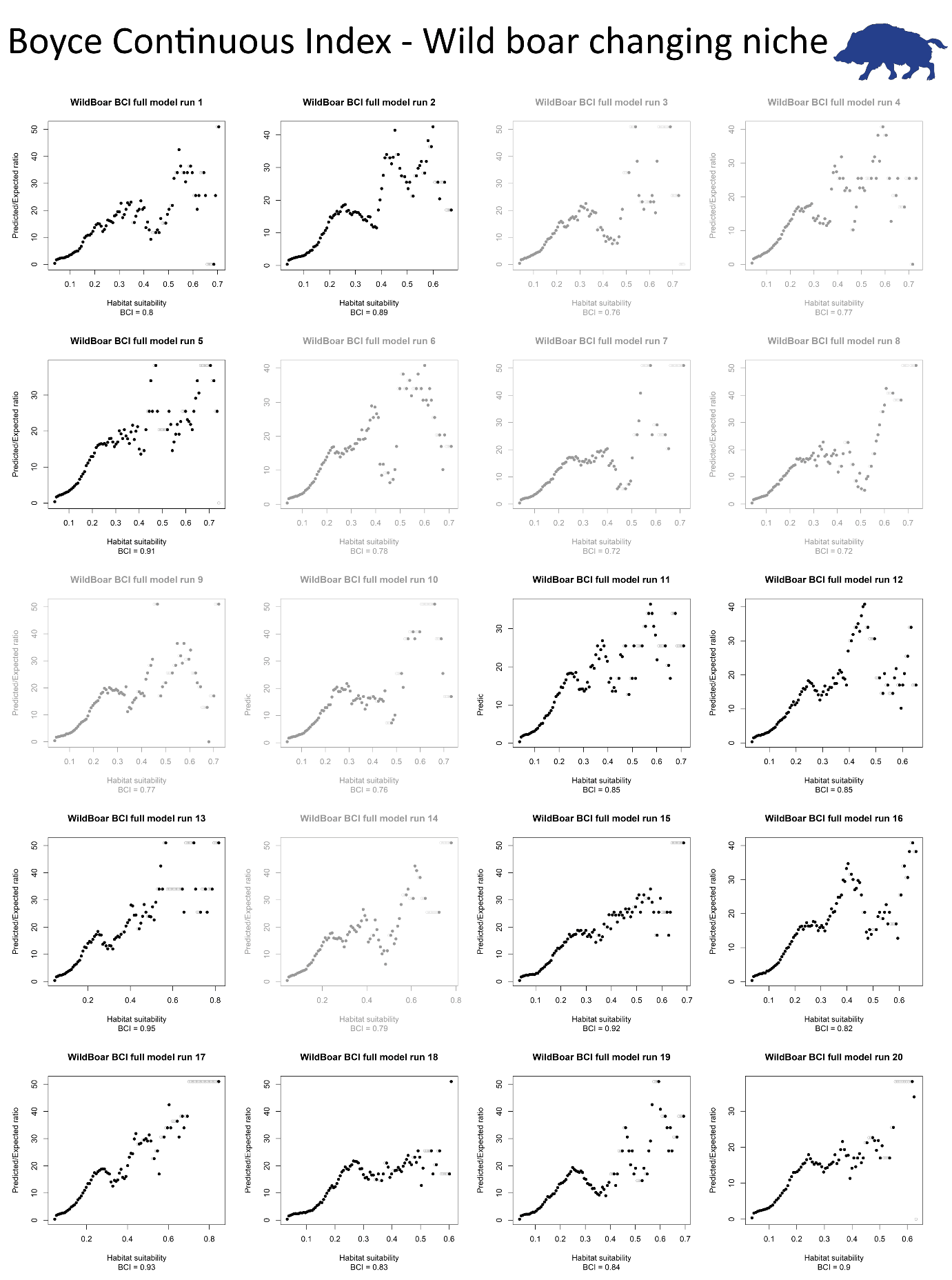

Figure S 9: Boyce Continuous Index (BCI) plots for each run of the changing niche model (= full model) in wild boar. In gray, runs with a BCI below the acceptance threshold (0.8)

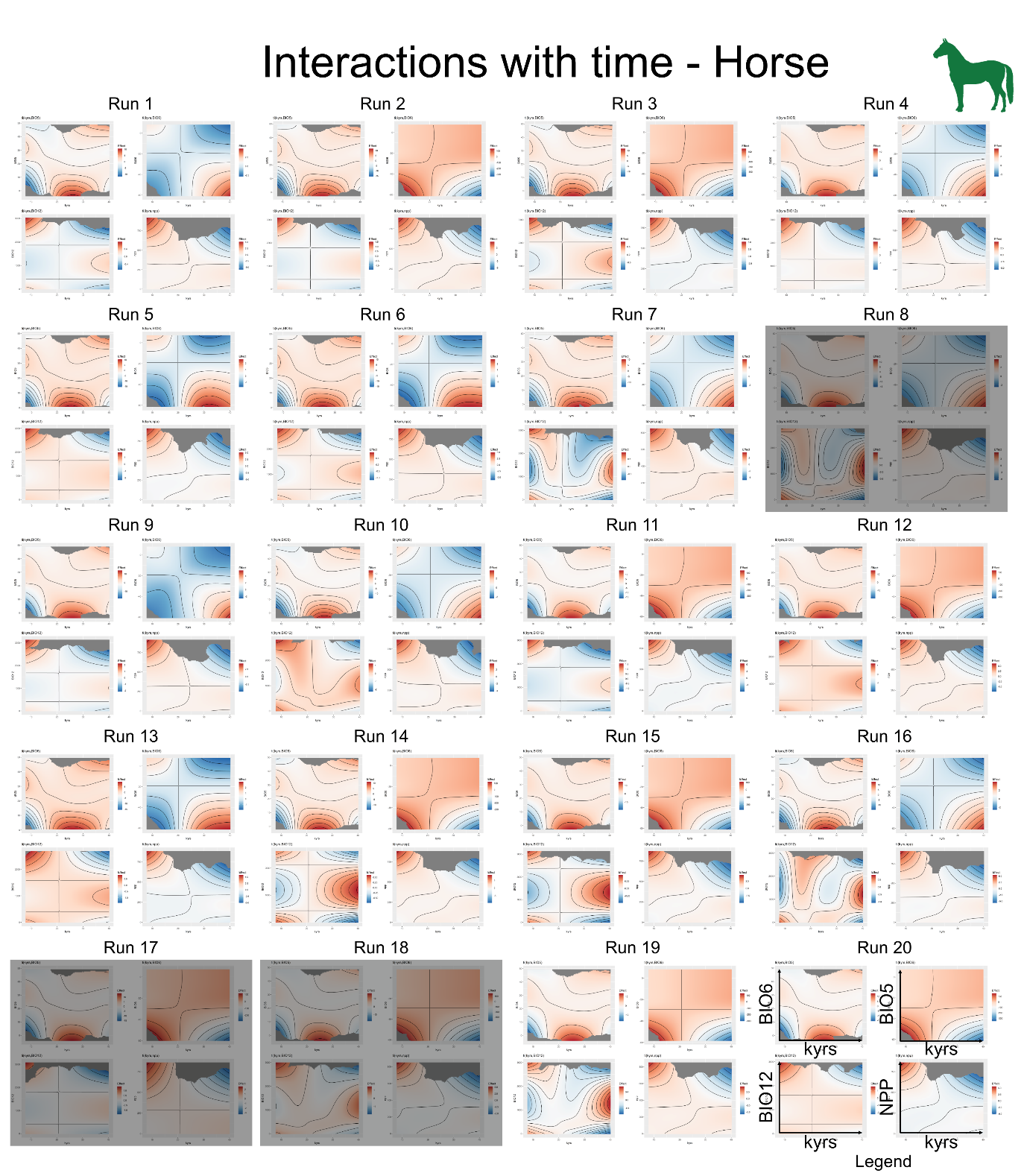

Figure S 10: Interactions with time for each run of the full model in horses. In gray, runs with a BCI below the acceptance threshold (0.8)

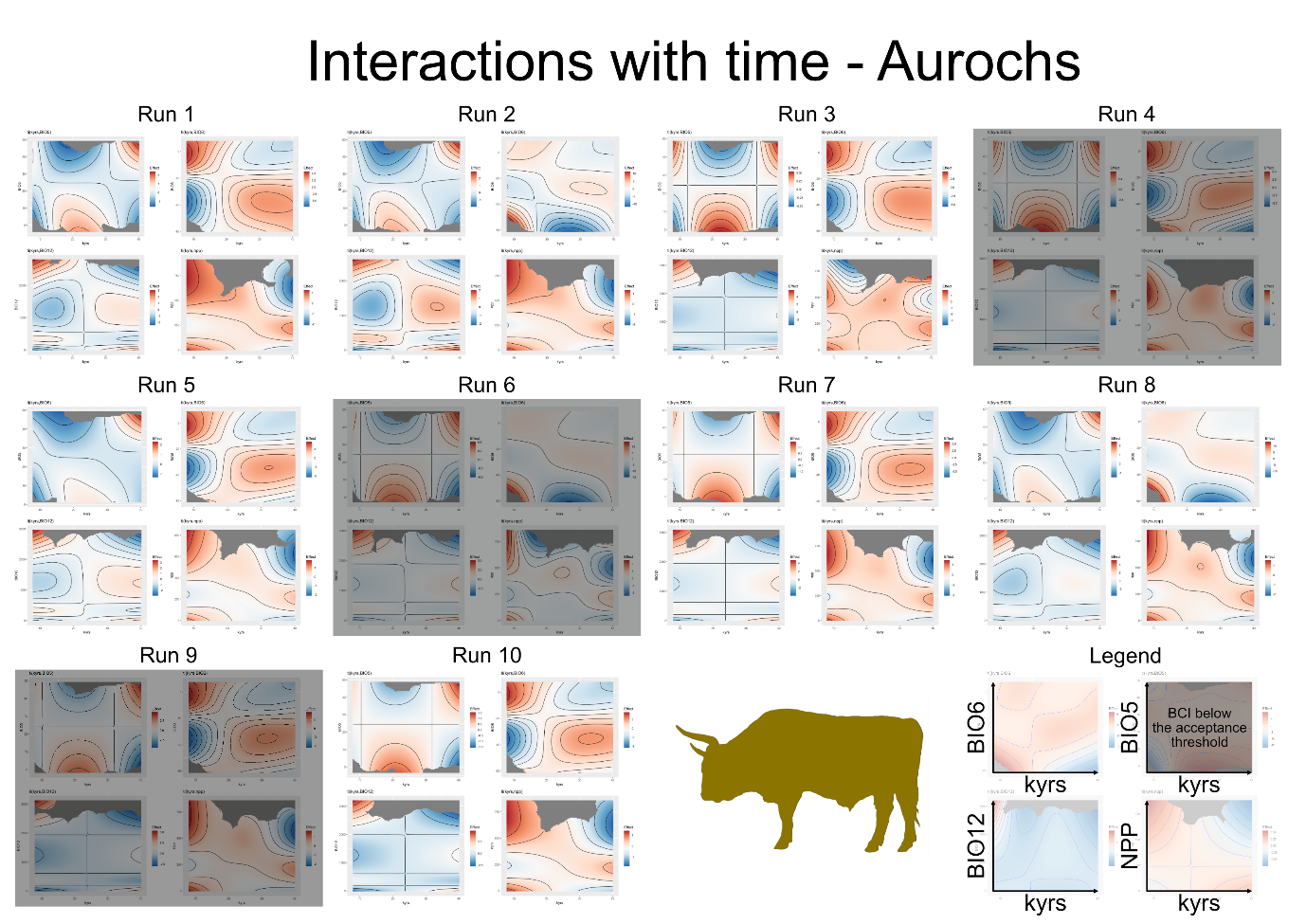

Figure S 11: Interactions with time for each run of the full model in aurochs. In gray, runs with a BCI below the acceptance threshold (0.8)

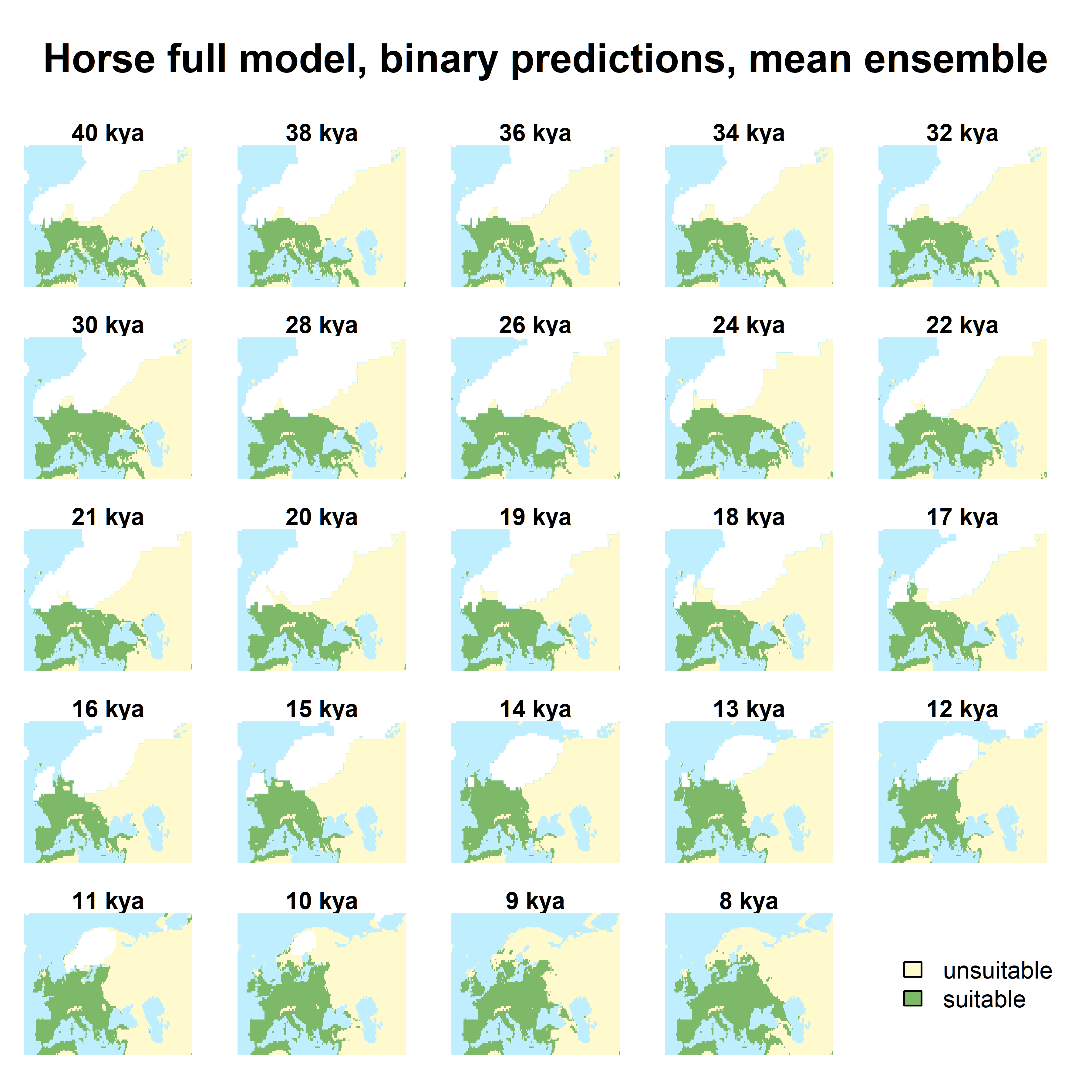

Figure S 12: Projection of the potential distribution of the horse based on the mean of the full ensemble.

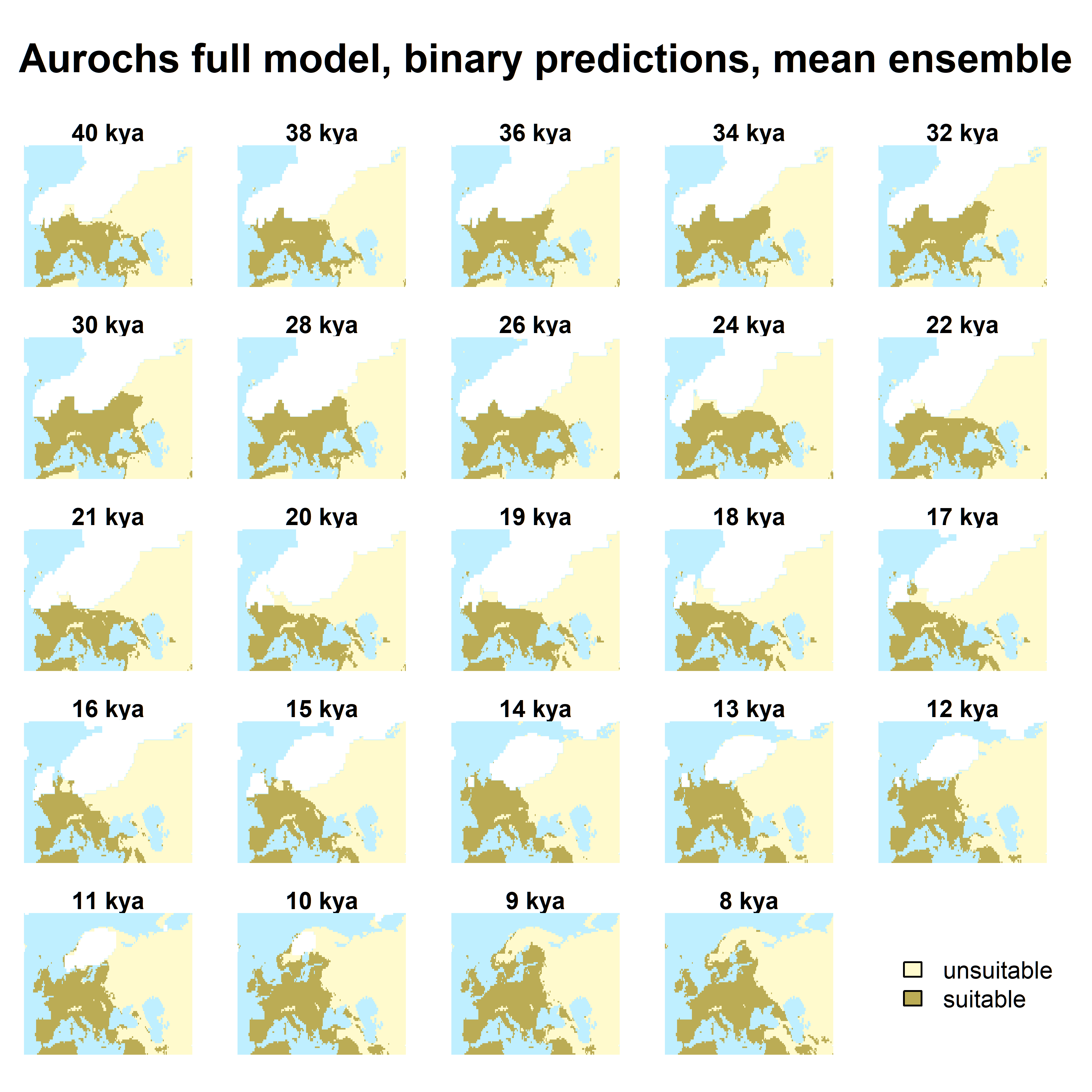

Figure S 13: Projection of the potential distribution of the aurochs based on the median of the full ensemble.

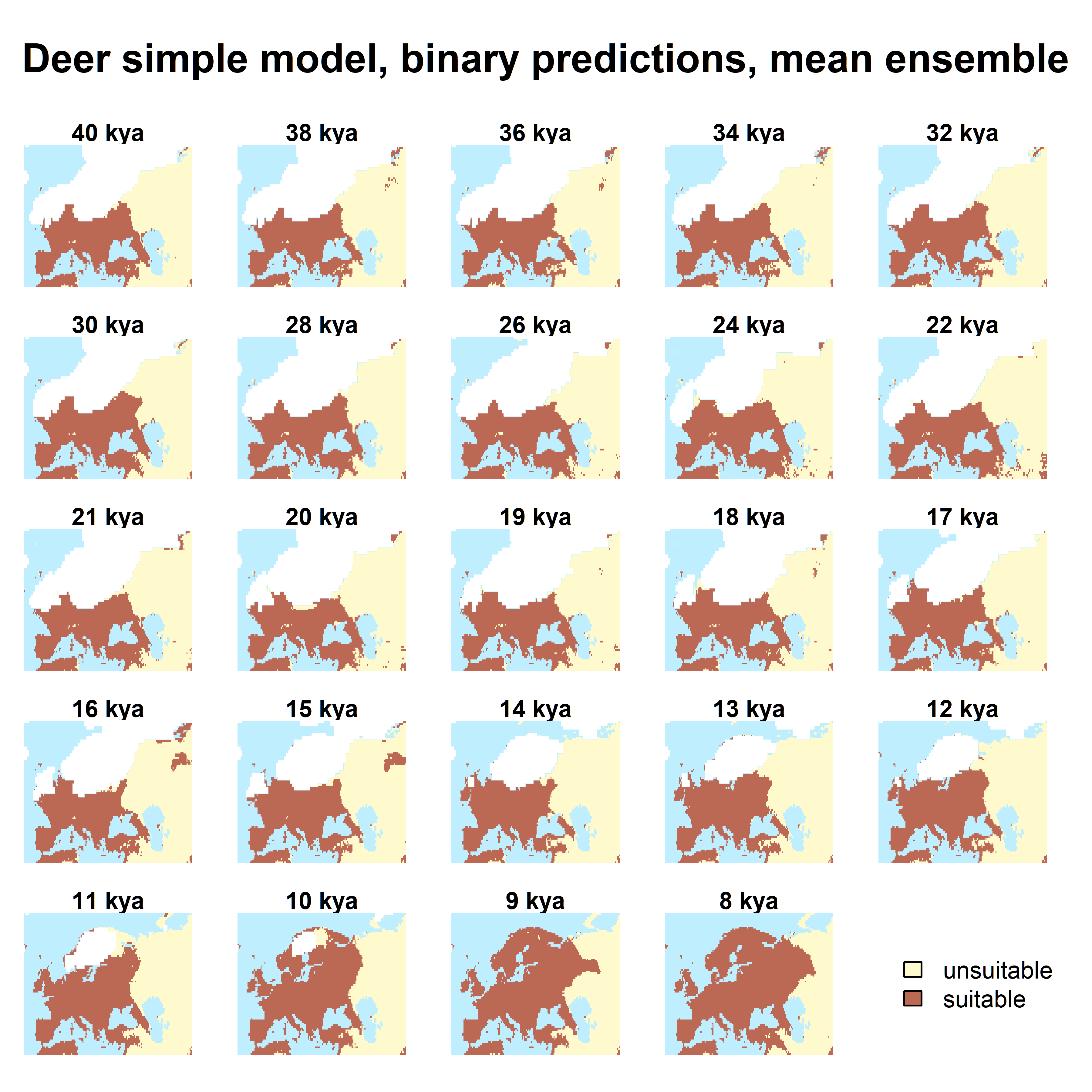

Figure S 14: Projection of the potential distribution of the dear based on the mean of the full ensemble.

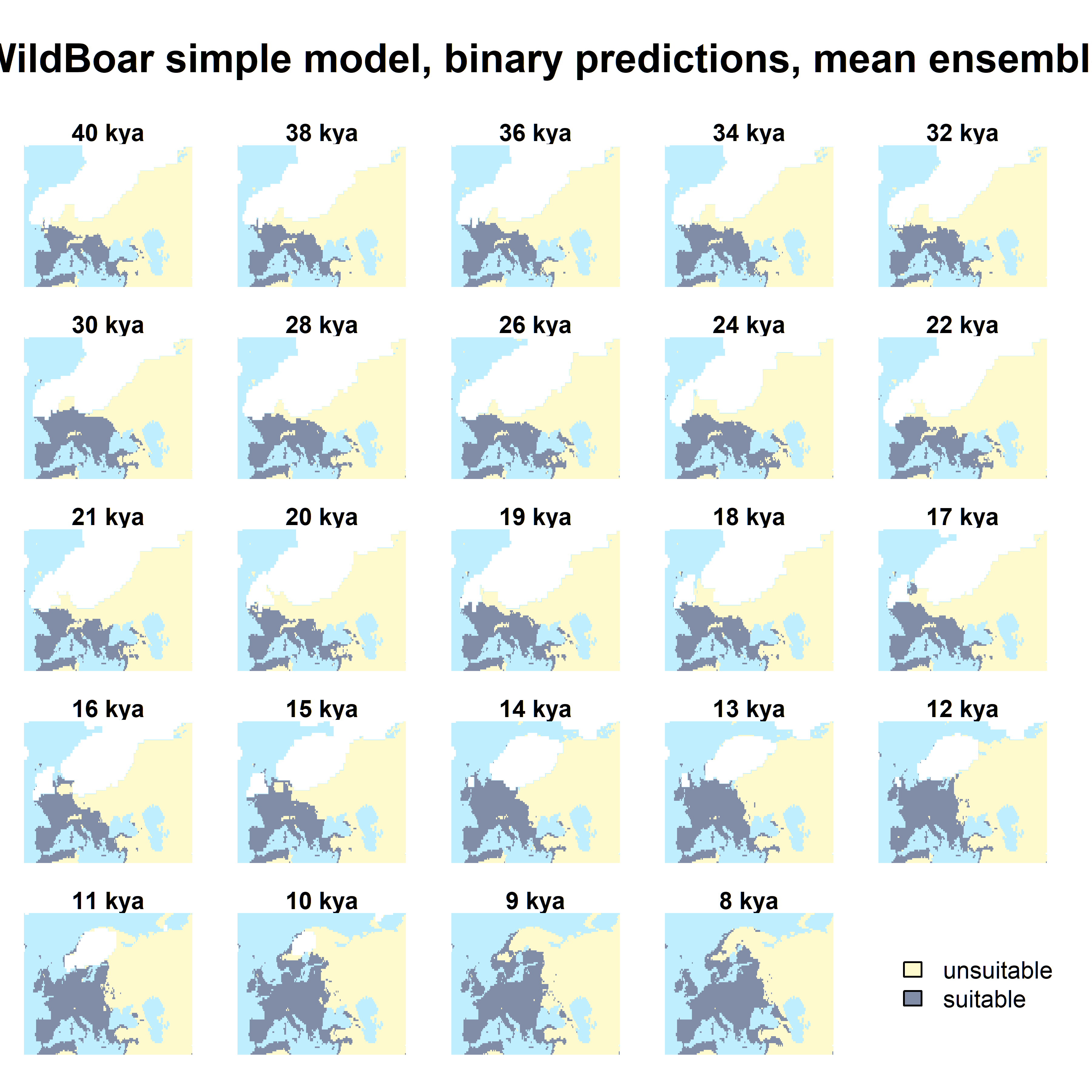

Figure S 15: Projection of the potential distribution of the wild boar based on the mean of the simple ensemble.

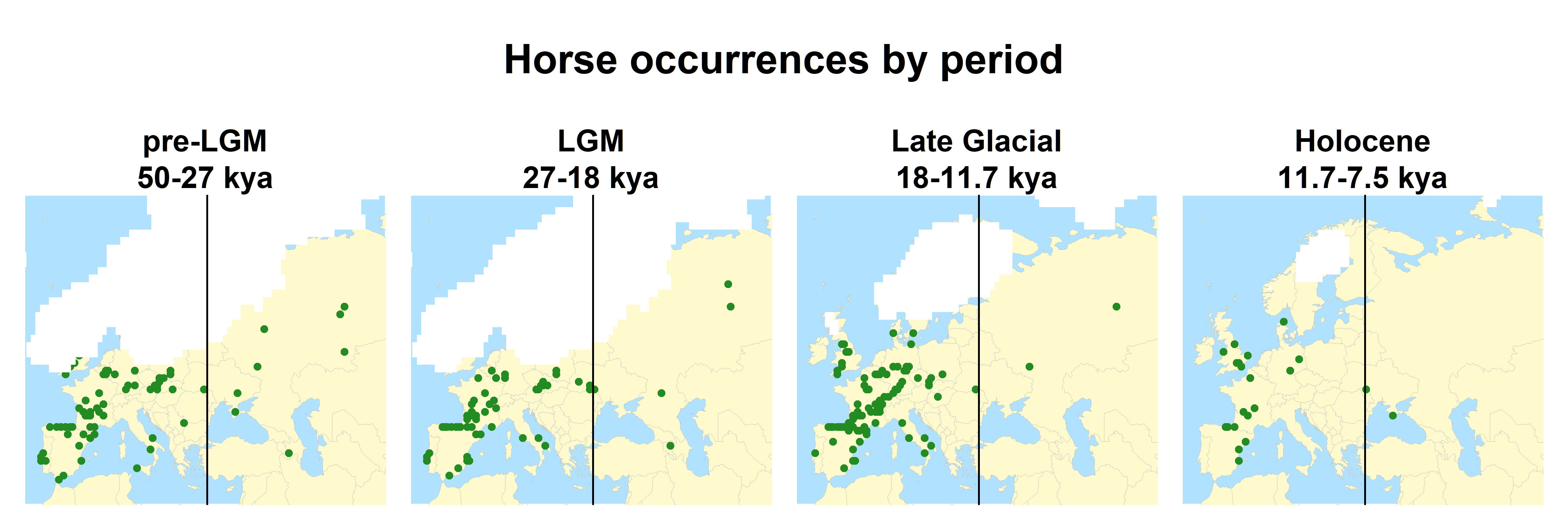

Figure S 16: Horse occurrences over the whole European continent

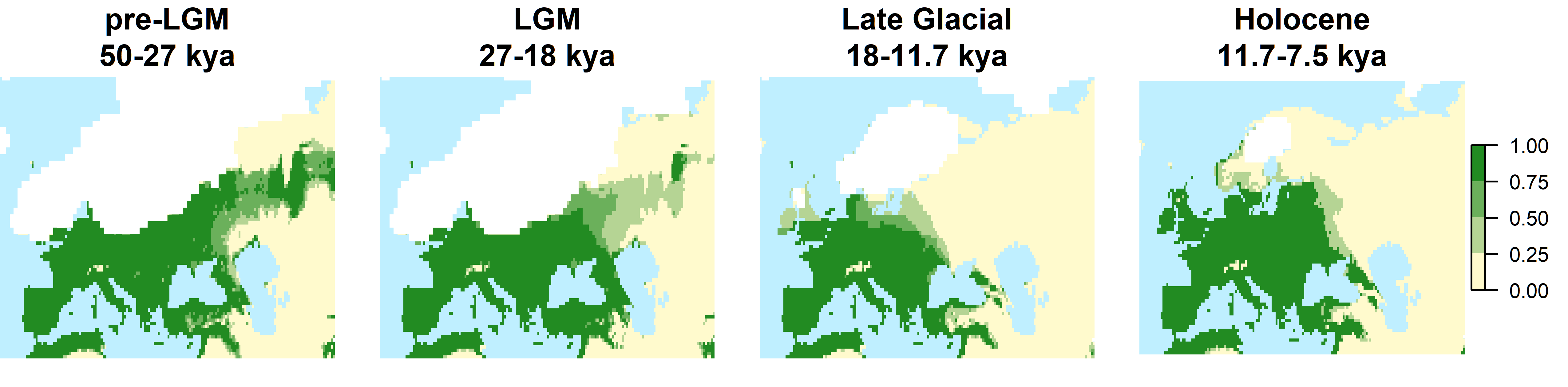

Figure S 17 Projection of the potential distribution of the horse based on the whole European dataset. Each frame has been obtained by averaging binary projections over individual time slices.
